# Occurrence of Aneuploidy Across the Range of Coast Redwood (*Sequoia sempervirens*)

**DOI:** 10.1101/2024.11.19.624336

**Authors:** Alexandra Sasha Nikolaeva, James Santangelo, Lydia Smith, Richard Dodd, Rasmus Nielsen

## Abstract

Aneuploidy, a condition characterized by an abnormal number of chromosomes, can have significant consequences for fitness of an organism, often manifesting in reduced fertility and other developmental challenges. In plants, aneuploidy is particularly complex to study, especially in polyploid species such as coast redwood (*Sequoia sempervirens*), which is a hexaploid conifer (2n=6x=66). This study leverages a novel Markov Chain Monte Carlo (MCMC) method based on sequence depth to investigate the occurrence of aneuploidy across the range of coast redwood.

We show that aneuploidy is prevalent in second-growth redwoods, predominantly as additional chromosomes, while tissue culture plants frequently experience chromosome loss. Although our study does not directly assess the fitness of aneuploids, the frequency of chromosomal instability observed in tissue culture plants compared to second-growth and old-growth trees raises questions about their long-term developmental viability and potential to become established trees. These findings have significant implications for redwood conservation and restoration strategies, especially as tissue culture becomes the primary mode of producing nursery stock plants used in reforestation.

## 2 Introduction

Aneuploidy—or a change in the base number (*n*) of homeologous choromosomes [1]—is generally considered deleterious, as an unbalanced chromosome dosage can lead to the loss of fitness of an organism. In many diploid organisms, including humans, aneuploidy is linked to a shorter life span and can cause certain health symptoms. For example, trisomy 21 is a genetic condition responsible for Down syndrome in humans and often associated with learning disabilities, congenital heart diseases, Alzheimer’s diseases, leukemia, cancers, etc [2].

While the causes and consequences of aneuploidy have been reasonably well-studied in humans and other animals, less research has focused on plants [3]. Some plants appear to tolerate aneuploidy better than others, especially polyploid plants that have not undergone the diploidization process [4]. For example, in hexaploid wheat (*Triticum aestivum*), changes in the copy number of a homeologous set of chromosomes (an extra or a missing chromosome 4B and 7A) did not significantly affect the expression levels of that set as a whole [5]. In contrast, in a population of synthetic tetraploid *Arabidopsis thaliana*, the presence of an extra chromosome 5 led to significant detrimental effects, including altered gene expression on the trisomic chromosome, changes in gene expression on other chromosomes, and overall genome instability [6]. Laboratory experiments with self-pollinated allopolyploids like *Brassica napus* have shown that aneuploids are selected against, as gene expression becomes imbalanced [7].

Key causes of aneuploidy in better-studied diploid genomes include incorrect attachments between chromosomes and spindle fibers, which can go undetected by the cell’s checkpoint systems. Extra centrosomes can create abnormal spindle shapes, leading to mis-segregation during meiosis or mitosis [8]. Meiotic aneuploidy most often arises from nondisjunction during chromosome segregation in gametes due to errors like premature chromatid separation [9], while mitotic aneuploidy occurs in somatic cells due to spindle assembly failures, often influenced by chemicals that disrupt microtubule dynamics [10]. Age-related defects in chromosome cohesion also contribute, as do stresses on DNA replication and repair processes. Factors like oxidative stress and mechanical stress during cell division further disrupt chromosome segregation [1].

Environmental factors are known to influence levels of aneuploidy in natural populations. On a cellular level, for example, water stress has been demonstrated to cause meiotic chromosome abnormalities in rice, barley, and other agricultural crops, often reducing fertility of male plants [9]. Given this effect, it is therefore not surprising that at the population level aneuploidy has been also found to occur at the extremes of non-agricultural species ranges at the edges of preferred climatic conditions, where the species is subject to increased environmental pressures. For example, in diploid Scots pine (*Pinus silvestris*) at the edges of its natural range in Khakassia, [11] found a wide range of genomic and chromosomal rearrangements, including aneuploidy and mixoploidy, which they attributed to a combination of factors such as dry and nutrient-poor conditions typical of mountainous, gravelly-stepped landscapes, as well as reproductive isolation of the population.

Coast redwood (*Sequoia sempervirens* (D. Don) Endl.,) is a long-lived conifer restricted to the coastal fog belt from southern Oregon to central California. It reproduces mainly asexually, but seed reproduction also has an important role in population dynamics [12]. The species is hexaploid [13–15] and a tentative autopolyploid [16], although earlier studies indicate partial allopolyploidy [14]. The degree of sequence differentiation among all six copies of redwood chromosomes has not been extensively studied and the number of chromosomes per gamete (gametophyte) is also not known. In a distantly related species of cypress (*Cupressus sempervirens*), megagametophytes have been found to contain “an even and odd series of DNA contents: 1C, 2C, 3C, 4C, 5C etc., where C is the amount of DNA in the haploid genome” [17].

The examination of allozyme inheritance patterns conducted by Rogers [18] indicates that hexasomic inheritance, in which each combination of homologous chromosomes is equally likely to be formed in a gamete, is largely preserved in redwood. These results exclude strictly disomic inheritance, where homologous chromosomes always pair in a particular configuration, making some combinations impossible. However, multisomic inheritance does not rule out the possibility of bivalent pairing configurations in an autopolyploids [3, 19], and that might also be true for coast redwood. A later study, examining the chromosome segregation patterns in meiosis [20] showed that many of the redwood chromosomes do pair as bivalents, although full multivalent meiotic rings and other chromosomal configurations, including monovalents, trivalents, and tetravalents were also present. It is possible that certain redwood chromosomes have started to diverge in bivalent pairs, but this process is slow and still allows for multivalent pairing.

The inconsistent meiotic behavior could help explain low (up to 15%) seed germination in coast redwood [21]. As Stebbins pointed out in a seminal paper on polyploid plants, [22], irregular chromosomal configurations often lead to sterility and formation of aneuploids. Later studies [23] suggested that formation of univalents and trivalents is often responsible for low fertility as such gametes often result in an unbalanced number of chromosomes in a cell. More recent studies confirmed this suggestion in *Brassica napus* plants, in which seed yield and pollen viability were inversely correlated with increasing aneuploidy [7]. Interestingly, seed viability in redwood tends to increase with age of the parent trees with the maximum viability reached when trees are over 250 years, before tapering off and decreasing at over 1,200 years of age, with many exceptions [24]. The extent to which seed viability is affected by imperfect segregation in meiosis and the resulting aneuploidy remains an open question.

In this research, we evaluated cases of aneuploidy across the geographic range and temporal scale of redwoods. We utilized short-read sequencing to identify aneuploidy in both old growth and second growth trees and compared these patterns with those observed in tissue-culture plants. The primary challenge in this research lies in accommodating technical variance in sequencing depth among individuals and chromosomes. While this is relatively straightforward and standard practice in organisms with well-described and curated genomes, such as humans and model organisms, it becomes significantly more complex in poorly understood hexaploid genomes. To address these challenges, we developed a new Markov Chain Monte Carlo (MCMC) method for inferring variation in chromosome number from population samples of non-model organisms. This approach allows us to accurately detect and analyze aneuploidy despite the inherent difficulties posed by the unique and understudied coast redwood genome.

## 3 Materials and Methods

### 3.1 Sample collection

To investigate aneuploidy across the range of coast redwood, we used a paired study design, in which pairs of populations are selected in such a way that the populations are geographically close but experience substantially different selective environments [25].

Sampling was conducted in unmanaged old-growth and redwood second-growth forests defined using the LEMMA forest structure dataset provided by the Save the Redwoods League [26]. The history of how the sampling site was managed was verified through consultations with landowners. Although we did not have an accurate estimate of the age of the trees – it is a notoriously difficult task to estimate the age of redwoods [27] – we assessed based on management history that the old-growth forests were over 300 years old (conservative estimate) and the second-growth trees were between 40-100 years old.

For each location, we collected between one and five foliage samples. The trees were located at a minimum of 60 meters from each other as this was the maximum distance between the clones reported in the most recent study on redwood clonality [28] to minimize our chances of sampling clonal trees in one location. The foliage was collected from the lowest branches of established trees or epicormic or basal sprouts when the lowest branches were not reachable, which was often the case in old-growth stands. In some locations, the so-called “sun foliage,” or the foliage that typically grows near the tree tops and has different phenotypic characteristics, was also collected from the ground. The location of each sample was recorded on Avenza maps (Avenza Maps™ v5.1.1).

Samples then were placed on ice and transported to the University of California, Berkeley campus within two days. Upon arrival, they were immediately stored in a -80°C freezer. In total, samples from 305 trees were collected (Figure 1).

**Figure 1:**
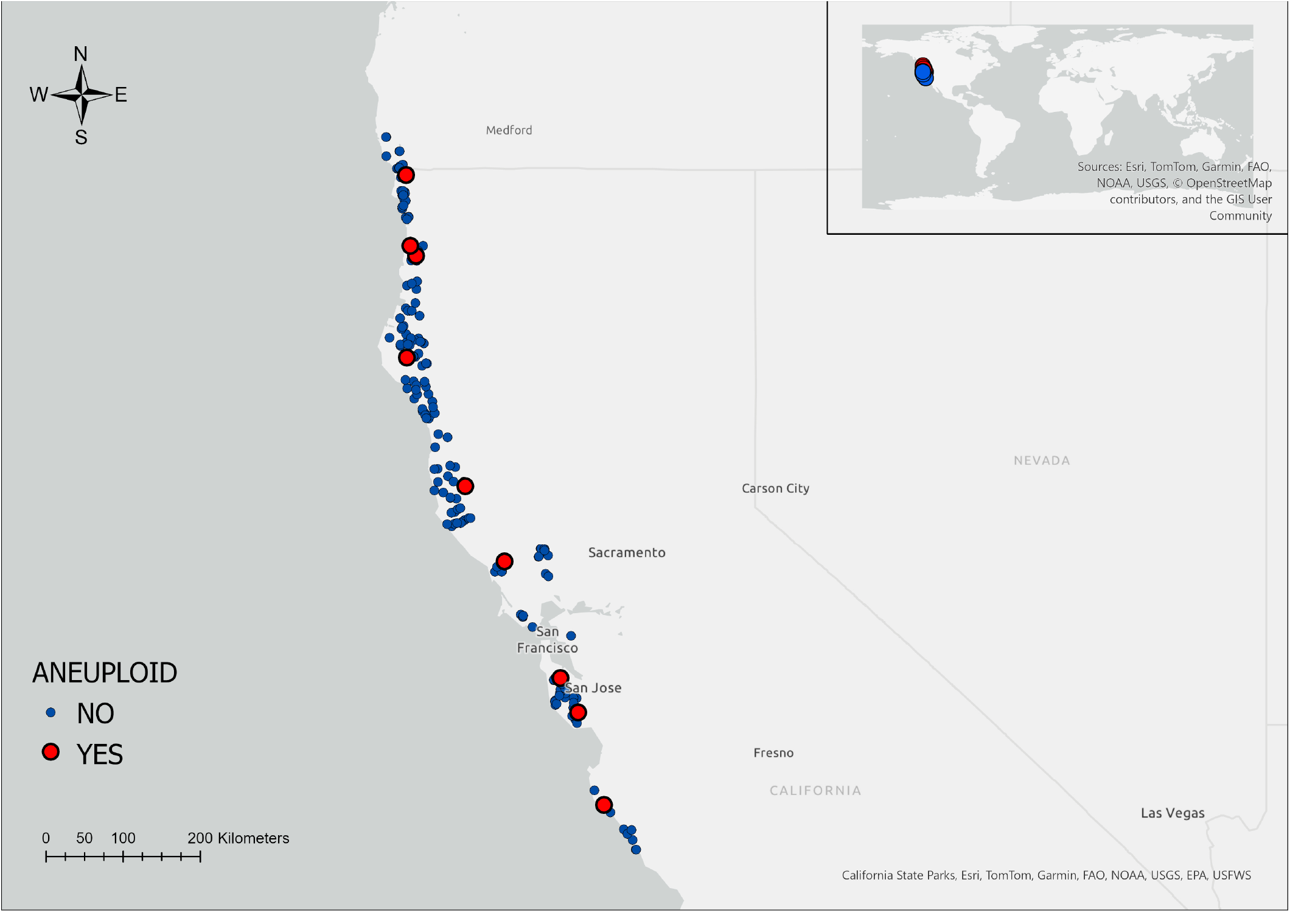
Sampling distribution across coast redwood range.

We also utilized a second coast redwood exome dataset described in [29]. The dataset included 82 samples, originating from tissue culture of plants collected as a part of the previous common garden experiment at the Russell Research Station [30]. The tissue culture from the common garden was propagated in a greenhouse and the foliage material was sequenced from the propagated cuttings at the University of California, Davis.

### 3.2 DNA extraction and sequencing

DNA was extracted from leaf tissue using a modified CTAB protocol [31] with changes made to the tissue preparation and DNA purification steps. Briefly, modifications to the protocol included a chloroform prewash applied to the homogenized tissue to remove the secondary metabolite compounds and a second ethanol wash. We also performed a magnetic bead clean-up using Solid Phase Reversible Immobilization (SPRI) Beads using a modified protocol from [32]. DNA was then resuspended in purified water. The concentration and purity of DNA were assessed on a SpectraMax M2 plate reader using the Biotium Accuclear High-Sensitivity Kit.

A portion of each extraction was diluted to 10 ng/μL at a volume of 110 μL using 10 mM Tris elution buffer, pH 8. The aliquot was sonicated on a qSonica Q800R sonicator at 40% amplitude and 15s on/off pulse for 5 minutes of active sonication time. A double-sided SPRI bead cleaning process was used to size-select fragmented DNA to ~300-500 bp and to concentrate to a volume of 12.5 μL. The enzymatic steps of library preparation followed a modified Kapa Hyper Prep (Roche Diagnostics) protocol. After end repair and a-tailing, a universal stub adapter was ligated, and then was extended to full length during amplification with TruSeq-style unique dual-indexing oligos provided by the Functional Genomics Laboratory (University of California, Berkeley). After the final cleaning, libraries were eluted in water. Samples were assessed for sizing on an agarose gel or Bioanalyzer DNA 1000 chip (Agilent Technologies) and quantified using the Biotium Accuclear High-Sensitivity Kit.

Libraries were then combined into 36 pools of 8 libraries per each pool. 1000 ng of library was used as input, such that each capture pool contained 8 *μ*g. Capture hybridizations were completed in sets of 4–8 captures at a time using the Twist Target Enrichment Standard Hybridization v2 Protocol and Kit. Manufacturer’s protocol was followed, except in addition to the standard blocking elements of the Twist kit, we also added additional adapter and indexing oligo blockers provided by Roche to compensate for the additional library material being used. (13.4 *μ*L of Universal Blocking Oligos and 50 *μ*L of Kapa Enhancer Reagent per capture reaction.) After captures, pools were split in half for enrichment PCR: the first half of the pool was amplified with 6–8 cycles of amplification depending on what number had worked well with previous captures. Then it was cleaned with SPRI beads and assessed on a Qubit v.2 Fluorometer. If the result was high, the cycle number was lowered for the second amplification reaction; if low, it was raised. After the second cleaning, all pools were combined and assessed on a Qubit. Final concentrations of the enriched capture pools ranged from 4.26–25.2 ng*/μ*L (median of 14.6 ng*/μ*L; average of 14.2 ng*/μ*L). At the Vincent J. Coates Genomics Sequencing Lab, all 36 captures were pooled together in equimolar amounts based on qPCR assessment using Kapa Biosystems Illumina standards (QB3 Genomics, UC Berkeley, Berkeley, CA, RRID:SCR 022170). This final pool of all captured libraries was sequenced across 4 lanes of Illumina NovaSeq X 10B to collect paired-end 150 bp data. (Sequencing was performed at the UCSF CAT, supported by UCSF PBBR, RRP IMIA, and NIH 1S10OD028511-01 grants.)

The total size of target space was 17.7Mb. Targets were selected using available annotations for redwood genome [33], and filtered to 60 percent identity using sequences that had an alignment to the custom conifer database, NCBI’s Plant RefSeq, or UniProt database. This selection criterion was chosen to enrich for the most conserved sequences across the target exome.

Out of 305 collected samples, 285 were sequenced. 1 sample had insufficient sequencing depth and was excluded from analysis. 2 samples were technical duplicates that were also removed. We also excluded a number of samples (8) that came from a suspected clonal group of trees, for the total of 274 samples we used in the analysis.

### 3.3 Reference genomes

For this analysis, we used two existing reference genomes. The first, published by The Redwood Genome Reference Genome Project (RGP) [33], is approximately 26.5 gigabases, reflecting the triploid size of the genome. There is little synteny of the RGP assembly to the genome assembly of sister species of giant sequoia (*Sequoiadendron giganteum*) [34]. Biologically, this lack of synteny is highly unlikely, given that giant sequoia’s genome is syntenic with both metasequoia genome (*Metasequoia glyptostroboides*) [34] and Japanese cedar (*Cryptomeria japonica*). We assumed that the lack of synteny between giant sequoia and coast redwood genomes is likely caused by an imperfect genome assembly.

The second, known as the PacBio redwood genome, was sequenced in 2020 using 33-fold long HiFi reads and is publicly available [35]. We utilized both the primary assembly of the PacBio genome (48.5 Gbp), and the full 51 Gbp of assembled unitigs. It is important to note that both reference genomes were assembled with tools primarily designed for phasing diploid genomes, and neither of the genomes are haplotype-resolved or chromosome-level.The RGP genome was assembled using the MaSuRCA assembler [36] and the HiRise scaffolder [37], whereas the PacBio genome was assembled with the HiFiasm assembler [38].

### 3.4 Confirming ploidy of the reference genomes

Given the hexaploid nature of the genome and the possibility of some gene copies missing in the reference genomes, leading to incorrect estimations of gene and chromosome loss, it was necessary to confirm that the annotation sequences we used in designing exome probes were found in exactly six copies in the reference. To confirm that, we used the set of exome sequences (CDS) from the annotation of the RGP triploid reference as a query input to the BLAST program (blastn v.2.9.0-2) [39] against the PacBio hexaploid reference:

~~~
blastn -db ssempervirens.p_utg.fa \
       -query 55K.fa \
       -out blast_results.txt \
       -num_threads 10 \
       -outfmt “6 qseqid sseqid pident length mismatch gapopen qstart qend sstart send evalue bitscore
~~~

The BLAST output was then filtered using custom R-scripts (GITHUB). Briefly, the code is used to calculate the frequency of hits for each query sequence. We create a data frame, `query_frequencies`, which represents the distribution of hit frequencies for each query sequence in a BLAST analysis. It is created by counting how many times each unique hit frequency (number of hits per query sequence) occurs. For example, if a frequency of 2 hits is observed for 10 query sequences, `query_frequencies` will have a row with `Var1 = 2` and `Freq = 10`. Each frequency is then shown as a bar in a histogram plot.

### 3.5 Normalization of reads depth

Raw reads were aligned to the closely related but diploid species of giant sequoia (*Sequoiadendron giganteum*) reference genome [40] using BWA (v. 0.7.17-r1188). Giant sequoia’s genome has a high level of synteny to cryptomeria’s genome [34] and this gave us higher confidence in the quality of the giant sequoia’s assembly and therefore, the success of our approach. Alignments were then deduplicated, collated, and sorted with samtools (v.1.17) [41]. Reads with a minimum quality score of 60 (MQ *>*= 60) were selected, and the idxstats tool from samtools was used to calculate the count of mapped reads to each chromosome.

To visualize chromosome specific differences in sequencing dosage, we computed a read depth for each individual sample *i* for each chromosome *j*, standardized by the total read depth of the sample and the total read depth for the chromosome over all samples:

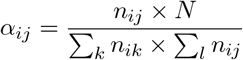

where *n*_*ij*_ is the read count for sample *i* on chromosome *j* and *N* = ∑^*i*^ *∑*^*j*^ *n*_*ij*_ is the total read count across all samples and chromosomes. A value of α_*ij*_ *>* 1 (or α_*ij*_ *<* 1) indicates that sample *i* has more (or fewer) reads on chromosome *j* than expected given the total number of reads for sample *i* and the total number of reads across all samples for chromosome *j*.

Taking into account the possibility of varying read depths due to differences in the number of amplification cycles, we stratified the dataset into five groups according to the total number of PCR cycles. Then α was calculated separately for each group.

### 3.6 Bayesian Model for determining ploidy per chromosome

We developed a Bayesian statistical framework to infer the number of chromosomes in coast redwood.

#### 3.6.1 Prior Distributions

The model assumes a uniform Dirichlet prior for the expected proportion of reads assigned to each chromosome: *λ* = (*λ*_1_, *λ*_2_, …, *λ*_*m*_), which can be interpreted as the length of the mapping target on each chromosome under the assumption of constant mapping probability along the length of the genome:

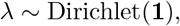

where **1** denotes a vector of ones, and *m* is the number of chromosomes.

The prior for the ploidy level (*k*_*ij*_) for individual *i* and chromosome *j*, is a reflected and truncated geometric prior distribution, centered around the expected ploidy of 6:

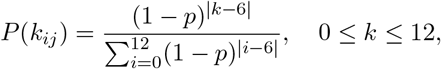

We assign a uniform[0, 1] hyperprior to *p*.

#### 3.6.2 Likelihood Function

The likelihood function is defined by a multinomial distribution, which models the read counts across different chromosomes for each individual as:

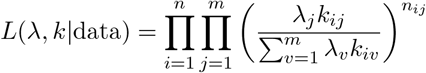

where *n* is total number of samples.

### 3.2 MCMC Algorithm

To estimate the posterior distributions of the parameters, we use a Metropolis-Hastings Markov Chain Monte Carlo (MCMC) approach. The MCMC algorithm iteratively updates the parameters *λ, k*, and *p* through a series of steps designed to explore the parameter space.

#### 3.7.1 Updating kernels

**Updates of** *λ* New values of *λ, λ*^*′*^, are proposed using a reflected exponential distribution, ensuring the new values remain within the permissible range, [0,1]. The update rule is given by:

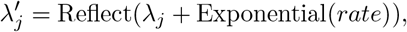

where the Reflect function iteratively sets *x ← −x* if *x <* 0 or *x ←* 2 *− x* if *x >* 1 until 0 ≤ *x* ≤ 1. This ensures that the proposed value of 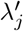 is between 0 and 1 and maintains symmetry of the update kernel. Other values of *λ* are then updated as

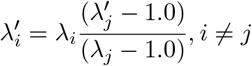

Because of the symmetric proposal kernel and the uniform Dirichlet prior, only the likehood enters into the Metropolis-Hastings ratio for this update. The value of the *rate* used in our analyses is 100.0 corresponding to an exponential with mean 0.01.

**Update of** *k* The ploidy levels, {*k*_*ij*_}, are updated independently for each chromosome and each individuals using a simple symmetric random walk, on a circle of integers {0, 1, …, 12} where 0 and 12 are connected states such that symmetry of the updates are preserved. The Metropolis-Hastings ratio then includes the likelihoods and the prior, but not the symmetric update probabilities.

**Update of** *p* The parameter *p* is updated using a reflected exponential, with mean 0.01 (rate 100,0), similar to the one used for updates of *λ*. Because *p* ∼ *U* [0, 1] and the proposal kernel is symmetric, only the likelihood appears in the Metropolis-Hastings ratio.

**MCMC runs** The algorithm cycles between updating all {*k*_*ij*_}, all {*λ*_*j*_}, and *p* and runs for a predefined number of iterations (100000), using the first 10000 as a burn-in. Convergence is assessed by running multiple chains with different starting point and evaluating the variance of the parameters across chains and within chains, and the auto-correlation of the sampled values across iterations (See Appendix B).

The validity of the implementation was tested by ensuring that the prior distribution was recovered as the posterior when running the program without data and by comparing the likelihood calculations to an independent implementation in Mathematica. A program written in C implementing the algorithm is available from Github.

## 4 Results

### 4.1 Ploidy of the reference genomes

Of the 207,167 total sequences blasted, 54,771 sequences were found in sets of 6 (328,626 qseqid hits) (see definition of a set in the Methods section), reflecting the hexaploid nature of the genome (Figure 2) in the PacBio full genome. After filtering for duplicates, the resulting number of sequences in the reference set was 39,840.

**Figure 2:**
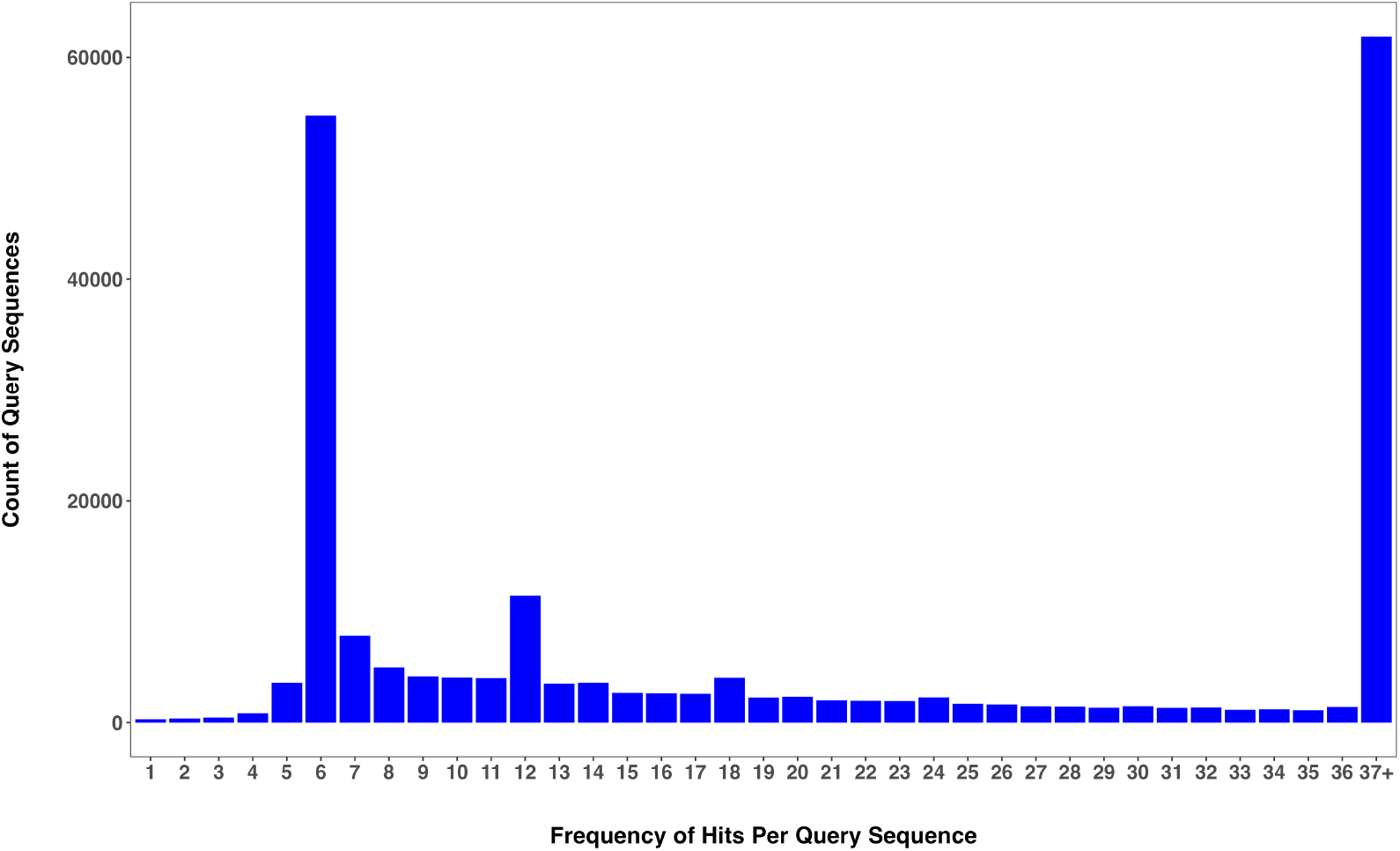
BLAST Query Frequency Distribution, PacBio reference genome.

However, most CDS sequences were found in high copy numbers, reaching up to 23,645 copies in one homologous set. This observation is consistent with previous findings that the coast redwood genome is rich in repetitive elements, which cover about 70% of the genome [33]. The average identity (pident value from the BLAST results table) between sequence copies found in sets of six was 99.36%, with a minimum value of 72.13% and a maximum value of 100%.

Another minor frequency peak was observed at 12 sequences per set, with 11,475 sets in total (Figure 2). From looking at the sequence identity of these sequences, it appeared that they could be split into groups of six—one group of six sequences among which the mean identity was comparable with the previous group (99.38%), and another group where the mean value of the pident was lower (95.63%). These sequences are possibly duplicates from an older event, either as individual gene duplicates or perhaps an older whole genome duplication event.

Repeating this analysis for the RGP genome, we found that most often CDS sequences were found in sets of four (22,105 unique sequences), followed by sets of three (19,638 sequences), indicating that this reference genome might not be triploid for all coding sequences, as expected (Figure 3). This finding disagrees with the results of the kmer analysis of the RGP genome using GenomeScope 2.0 software [42], which indicated triploidy of the reference.

**Figure 3:**
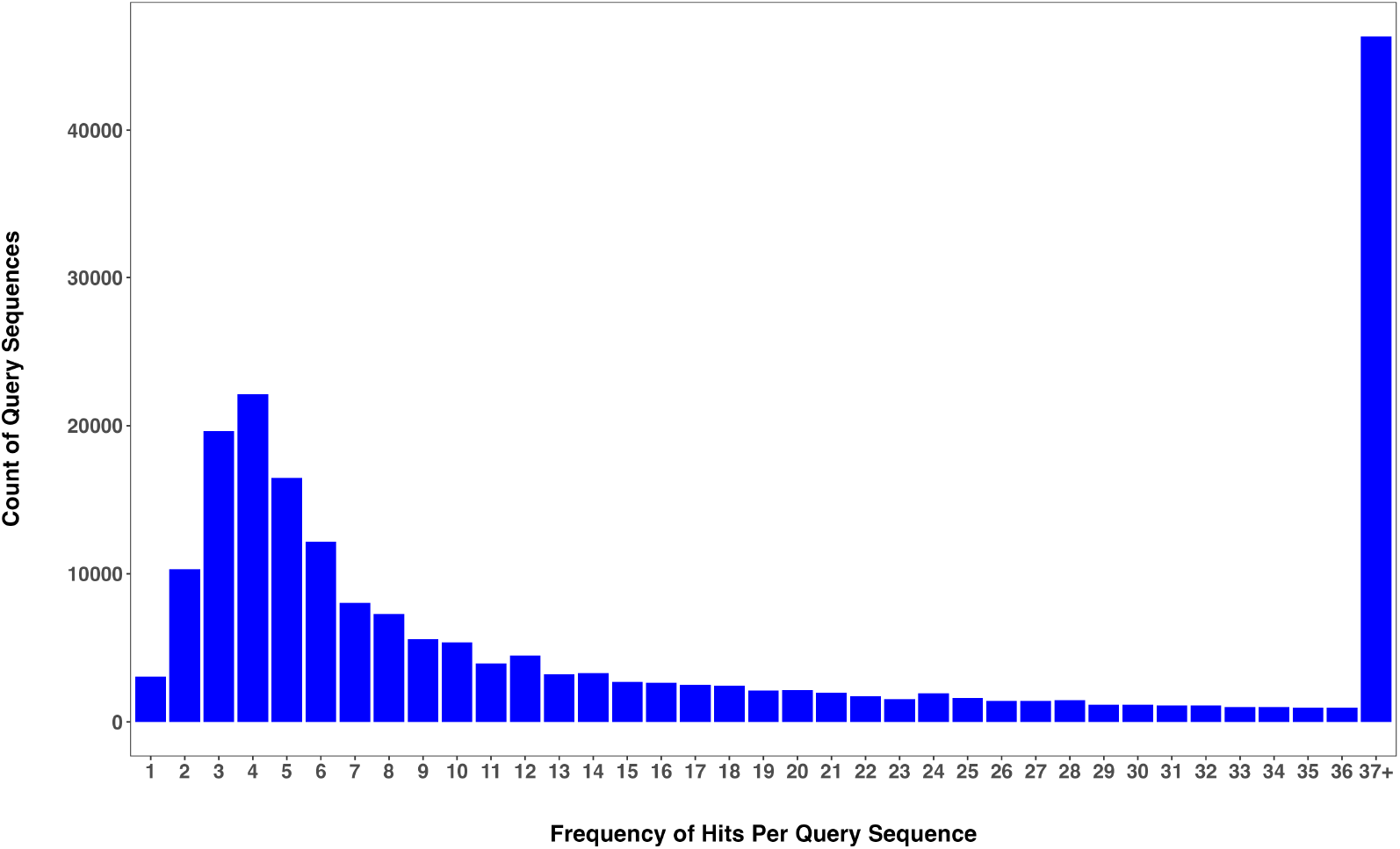
BLAST Query Frequency Distribution, RGP reference genome.

### 4.2 Ploidy per chromosome

The total number of reads (raw depth) aligning to giant sequoia (*Sequoiadendron giganteum*) chromosomes is shown in Figure 4, with the average of 9.13 × 10^2^ reads aligning per chromosome.

**Figure 4:**
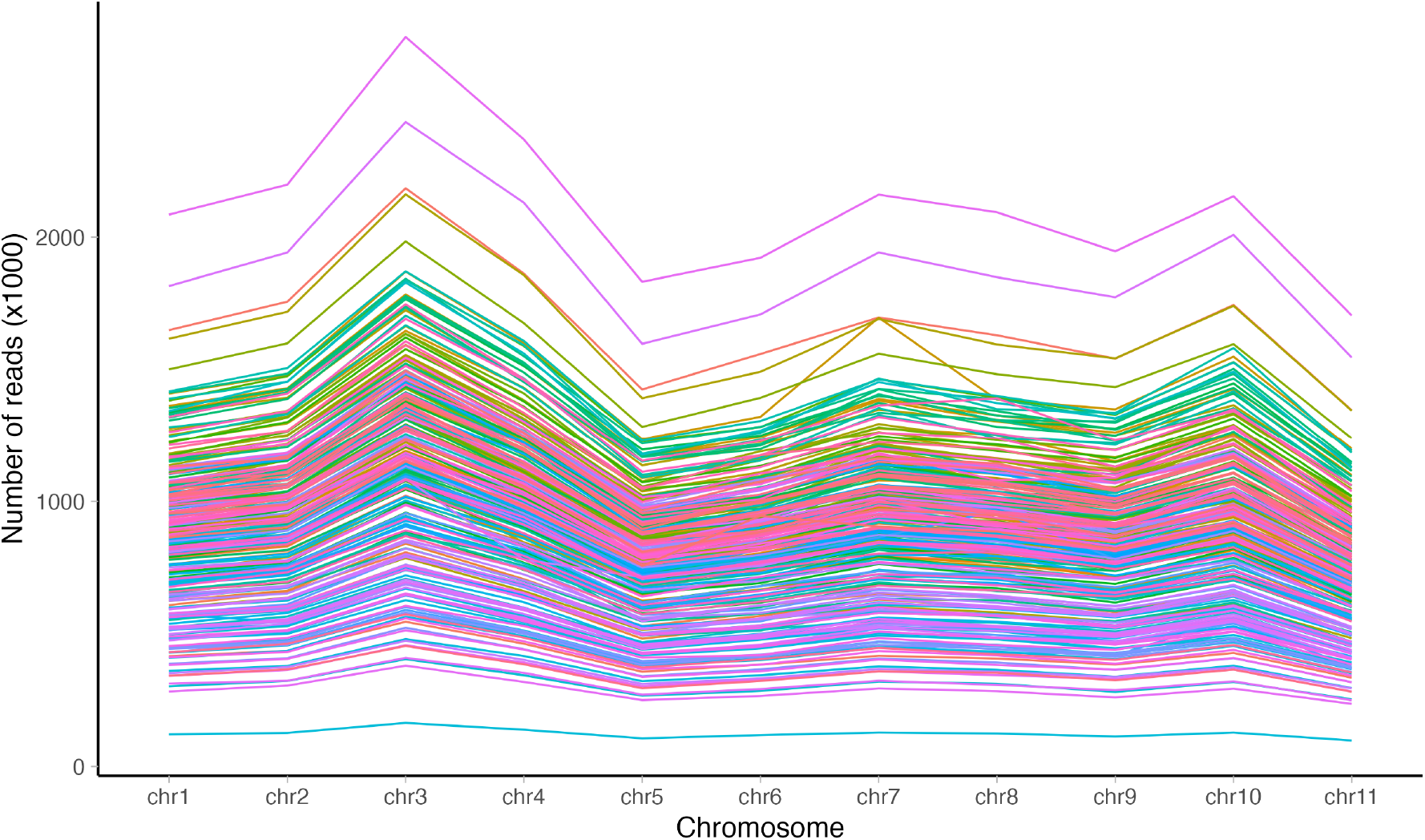
Raw read depth in coast redwood (*Sequoia sempervirens*) in relation to giant sequoia (*Sequoiadendron giganteum*) chromosomes. Each line represents an individual sample.

There were variations in *α* on all chromosomes except for chr2 and chr 5, characterized by either an increase or a decrease in *α* in the expected number of reads, indicating a deviation from the hexaploid chromosome number (Figure 5). To statistically assess these deviations and estimate the ploidy of each chromosome within each sample, we applied the MCMC algorithm described above.

**Figure 5:**
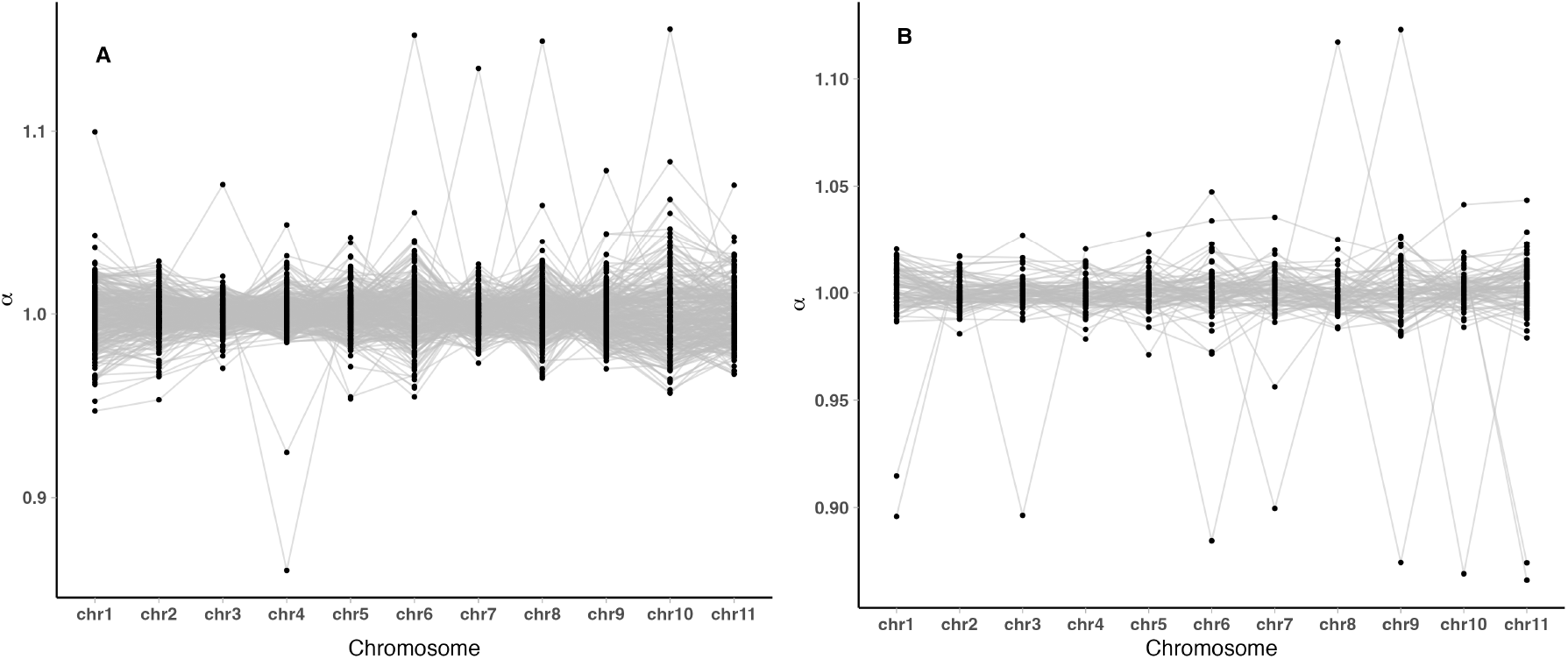
(A) Read depth normalization (*α*) in coast redwood (*Sequoia sempervirens*) in relation to giant sequoia (*Sequoiadendron giganteum*) chromosomes, this study; (B) Read depth normalization (*α*) in coast redwood (*Sequoia sempervirens*) in relation to giant sequoia (*Sequoiadendron giganteum*) chromosomes, as shown in the study by [29]. Each line represents an individual sample.

**Figure 6:**
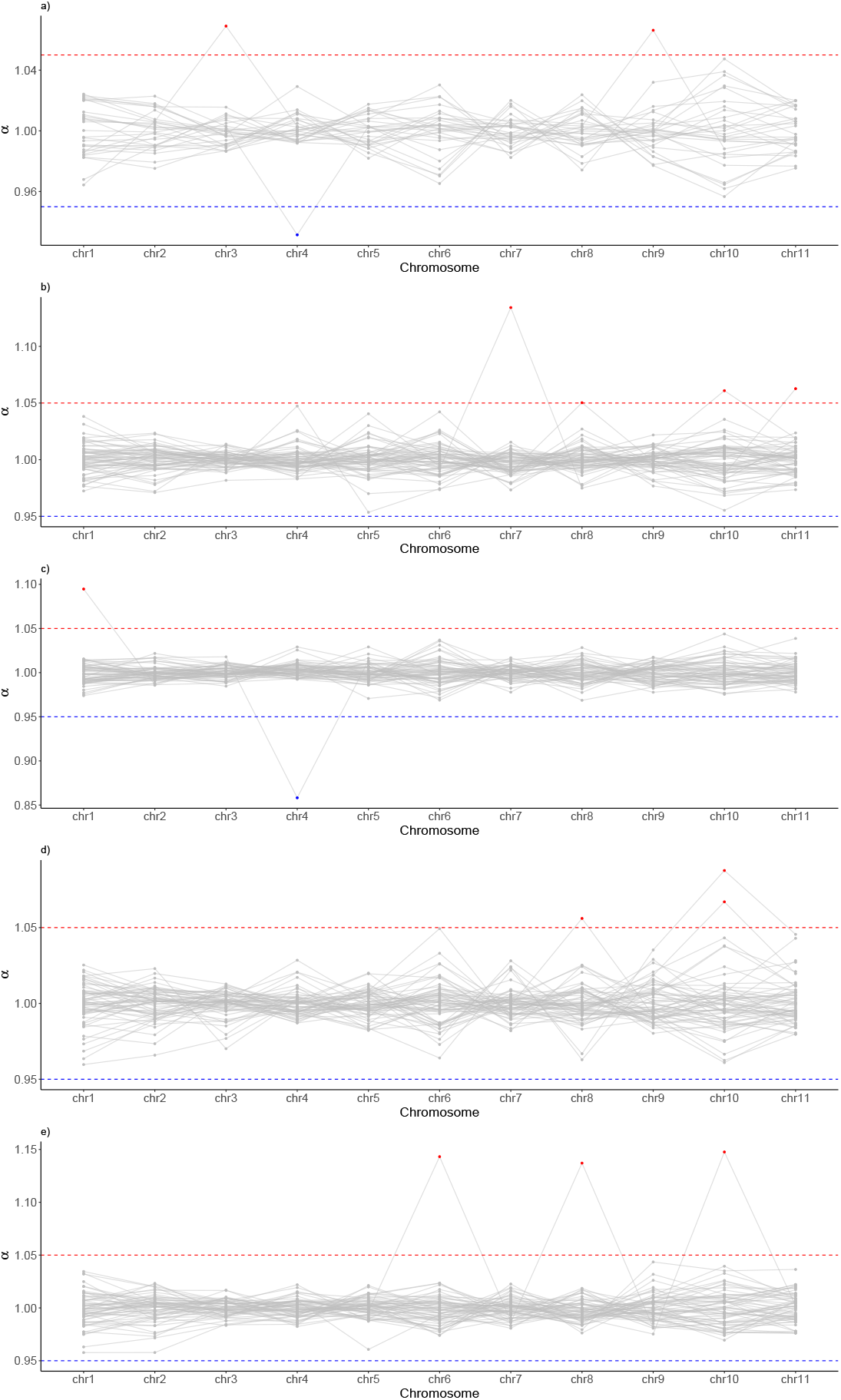
Read depth normalization by PCR group, a) -e) PCR groups from 1 to 5

**Figure 7:**
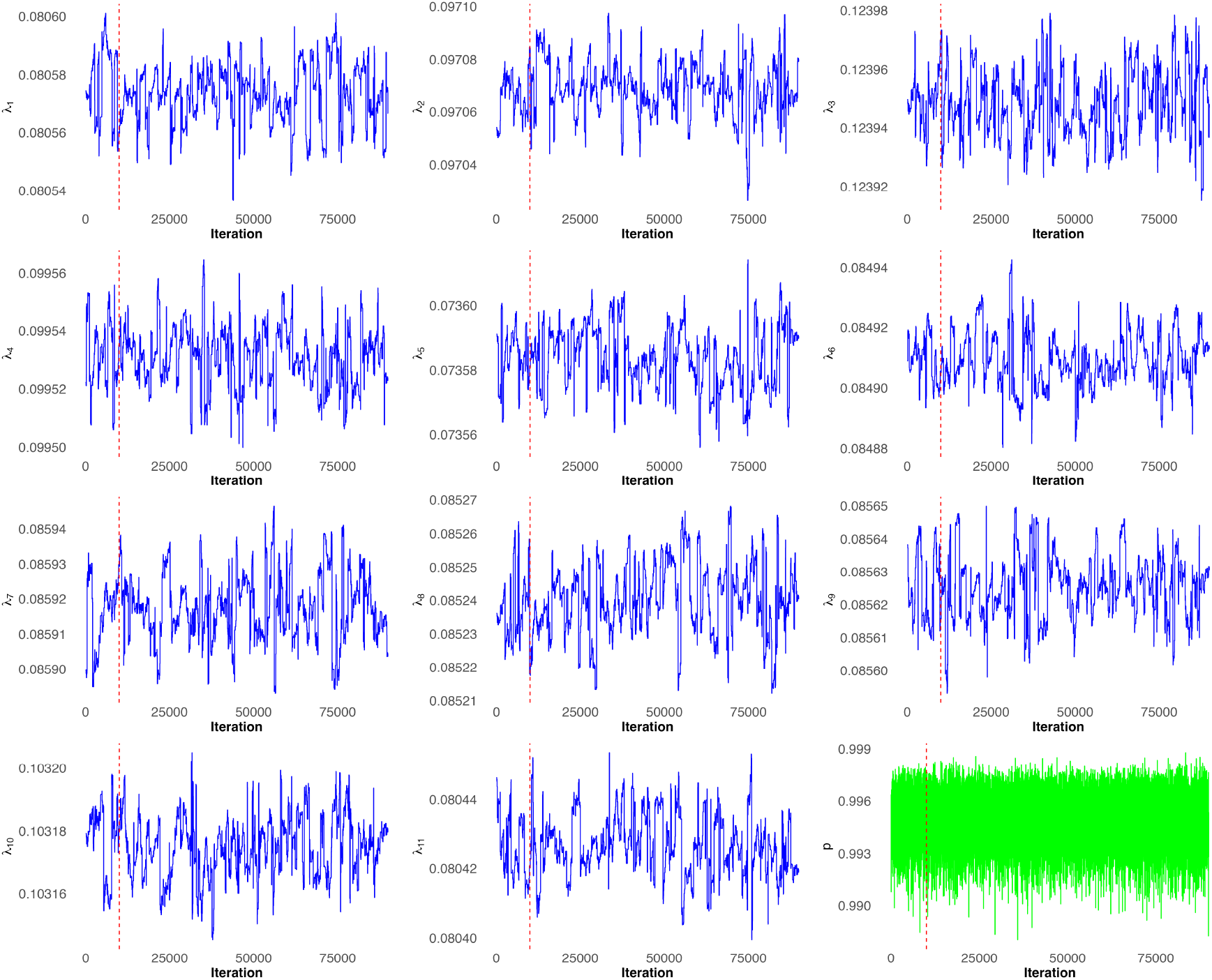
Trace plots for MCMC parameters

Additionally, to address the observed variance in *α*, we also ran the MCMC algorithm separately for each subgroup analysis based on the number of PCR amplification cycles for each sample. This allowed us to assess whether variations in amplification cycles might have influenced chromosome counts inferred by the MCMC. However, stratification by PCR cycle groups did not change the results.

In total, aneuploidy was detected in 9 samples (out of 274) and 8 of these samples showed an increase in the number of chromosomes per set, with a ploidy of 7, while one sample had a loss of a chromosome (Table 2).

**Table 1:** Aneuploidy counts across all samples from this study and the tissue culture dataset [29].

| Chromosome | This study (n = 274) |  | Tissue Culture (n = 82) |  |
| --- | --- | --- | --- | --- |
|  | Ploidy 5 | Ploidy 7 | Ploidy 5 | Ploidy 7 |
| Chr1 | 0 | 1 | 2 | 0 |
| Chr2 | 0 | 0 | 0 | 0 |
| Chr3 | 0 | 1 | 1 | 0 |
| Chr4 | 1 | 0 | 0 | 0 |
| Chr5 | 0 | 0 | 0 | 0 |
| Chr6 | 0 | 1 | 1 | 0 |
| Chr7 | 0 | 1 | 1 | 0 |
| Chr8 | 0 | 1 | 0 | 1 |
| Chr9 | 0 | 1 | 1 | 1 |
| Chr10 | 0 | 2 | 1 | 0 |
| Chr11 | 0 | 0 | 2 | 0 |

**Table 2:** Aneuploidy counts by growth stage across all samples from this study and the tissue culture dataset [29].

|  | Old growth | Second growth | Tissue culture |
| --- | --- | --- | --- |
| **Loss of chromosome** | 0 | 1 | 9 |
| **Gain of chromosome** | 0 | 8 | 2 |
| **Total aneuploids** | 0 | 9 | 11 |
| **No aneuploidy** | 56 | 209 | 71 |
| **Total trees** | 56 | 218 | 82 |
| **Percent aneuploids** | 0 | 4.3062 | 13.4146 |

An increase on chromosome 10 was observed most frequently, with 2 samples affected. Other chromosomes with increases in ploidy number were 1, 3, 6, 7, 8 and 9. There was also a loss of chromosome for 1 sample on chromosome 4.

In the tissue-culture dataset, we observed 11 samples (of 82) with aneuploidy present. Two of the samples had an increase of the chromosomal number to 7 (on chromosomes 8 and 9), and the rest had a loss of chromosome (chromosomes 1, 3, 6, 7, 9, 10 and 11).

The number of samples with an increase in the chromosomal number per set (ploidy of 7) was comparable between the two datasets (Fisher’s exact test *p* ≈ 0.535). The number of samples with a decrease or loss of chromosome is significantly higher in the tissue-culture dataset (Fisher’s exact test *p* ≈ 1.039 *×* 10^*−*5^).

### 4.3 Temporal and geographic distribution of aneuploid trees

The GPS coordinates were recorded for every tree in the dataset. Instances of aneuploidy were found throughout the range of species (Figure 1) and there was no geographic clustering of aneuploids. The only sample with a missing chromosome (ploidy of 5) was found in the southern part.

The stand age data were also collected in the field. There were 56 old-growth trees and 218 second-growth trees. Aneuploidy was present in second-growth trees (4.3%) and in the tissue-culture dataset (13.41%) (Table 2), but missing from the old-growth trees. However, there was no statistically significant difference between the number of aneuploids in old-growth and second-growth trees. (one-tailed Fisher’s exact test *P* ≈ 0.21).

We were not able to identify the exact geographic locations of the samples in the tissue-culture dataset, although the sampling distribution was range-wide. All of the samples from the second dataset were seedlings grown from the tissue culture [29].

## 5 Discussion

### 5.1 Detection of aneuploidy and study limitations

Recent studies investigating aneuploidy events in plants have found that aneuploids are typically located at the edges of a species’ range, signaling a potential stress-related genome restructuring that could also be adaptive [11]. However, aneuploidy in plants remains poorly studied due to the lack of high-quality reference genomes. Polyploid plants and autopolyploids present additional challenges since detecting such chromosomal variation often involves extensive laboratory techniques such as measuring DNA content via flow cytometry [43] or QF-PCR [44].

In this study, we explored chromosome numbers in coast redwood populations across the range using a new computational method that utilizes sequencing depth data. However, this method might not be applicable to detecting partial aneuploids where only parts of chromosomes are added or missing. Mosaic aneuploidy, where only some of the cells might be aneuploid, would also be hard to detect using our method. Chimera trees with various levels of mosaicism have been recorded in coast redwood [45], and more studies are needed to confirm the frequency of such mosaicism in redwood populations.

Additionally, our assumption was that all trees in the two datasets were hexaploid, which means our method cannot distinguish between a hexaploid and a tetraploid plant with aneuploidy.

Aneuploidy can arise from errors during meiosis and mitosis [46]. When mechanisms ensuring correct segregation of chromosomes during these processes fail, the resulting cells might have an unbalanced number of chromosomes. Here we define aneuploidy as a whole-chromosome gain or loss. As more accurate chromosome-level reference genome for the species become available, it will open up more avenues for investigating other structural variations such as deletions, insertions, and translocations.

### 5.2 Reference set of CDS sequences

Our BLAST analysis is a relatively simple way to confirm the hexaploid nature of the genome. It is notable that the most frequent category in which CDS sequences were found was 6, but we also found sequences in sets of 12. These sequences are possibly sequences that existed in two copies in the original diploid genome that then underwent two rounds of whole genome duplication, resulting in 12 copies of that gene. It is possible that these sequences date back to the genome duplication event that occurred at the base of all seed plants [47]. The distribution of frequencies of CDS sequences in the gametophyte reference genome (RGP), however, did not show the expected pattern of 3 BLAST hits per sequence. Instead, we observed that sequences were most often present in copies of 4, followed by sets of 3. This could be explained biologically by the true presence of four copies of the genome in the gametophyte, but the more likely explanation is an imperfect genome assembly, where the assembler might have failed to collapse repetitive sequences properly, possibly due to a higher than expected differentiation between the homologous haplotypes. We also found that this disagrees with the results of the GenomeScope2.0 analysis that indicated the triploidy of the RGP reference. Similar disagreements have been previously reported for the baobab genome *Adansonia digitata* [48], where GenomeScope2.0 suggested a diploid homozygous genome for a confirmed tetraploid, aligning with the caution from the GenomeScope2.0 authors [42] that their tool may underestimate ploidy levels beyond certain heterozygosity thresholds. One possible reason could be that, at higher levels of heterozygosity, k-mers become too divergent to be consistently recognized as matching pairs, potentially leading to an underestimation of ploidy. However, the precise factors contributing to this limitation remain unclear.

### 5.3 Consequences of aneuploidy on tree fitness

Coast redwood is a long-lived species that employs a clonal mode of reproduction [28]. It is common for polyploids to reproduce vegetatively and such mode is often regarded as an escape strategy from the barriers to sexual reproduction [49]. This strategy might be successful for polyploids needing to occupy changing ecological niches during significant environmental perturbations [50] and do so quickly, but it might come at a cost to individual plants that harbor substantial and detrimental structural genome changes [51].

Clonality complicates our understanding of temporal dynamics in plant systems due to the lack of discrete generation times and the longevity of the species. Here, we refer to tree age as the number of years from seed germination to an established tree. Perhaps counterintuitively, the least mature from the individual plant age standpoint tissue-culture plants in our study might be biologically the oldest among the samples we analyzed, as they have not undergone meiotic recombination. In mammals, incidences of aneuploidy increase with maternal age [51], and similar effects have been recorded in plants [52] where the seed age was positively correlated with karyotypical instability. Therefore, our finding of a higher number of aneuploids in the tissue-culture plants might perhaps be best explained by the increased biological age of these samples.

In redwoods, there may be somatic mechanisms that limit the propagation of aneuploidy throughout the adult plant tissues. As adult redwoods continue to grow, such mechanisms might help discard aneuploid tissues, preventing their propagation. Newly generated euploid cells might outcompete aneuploid tissues and therefore not allow the aneuploid sections to become widespread.

Evidence from studies on clonal plants, such as aspen [53], suggests a localized accumulation of somatic mutations where there are lower rates of mutations present in the tissues that contribute to progeny versus those that do not. This indicates that certain mechanisms may selectively control somatic mutations within the organism, helping maintain genetic stability across tissue generations. However, when an aneuploid tissue-culture is propagated and if the ratio of the euploid to aneuploid cells is low, aneuploidy might persist throughout the tissue, giving rise to an aneuploid plant. Because these plants are not subject to competition or environmental pressures since they are often grown in greenhouse conditions, the detrimental effects of aneuploidy might not become immediately evident. However, the potential detrimental effects might be what might explain the reduction of aneuploids in wild populations.

The reduced environmental completion is what might also help aneuploid clones survive in the wild. In the clonal system like this, individual clones that lose or gain a chromosome, might still be able to survive by pulling the resources from the original plant’s root system. A good example of such host dependency is albino redwood sprouts, which are lacking green chlorophyll [54] and are unable to survive on their own. Such sprouts depend on the host tree for resources. But as the competition for growing space increases and seedlings that originated from seed become more established, the detrimental effects of aneuploidy start to play a more significant role, giving trees with a complete set of chromosome a competitive advantage. But such competition is not limited to clonal vs. seed trees, and it is likely that clones with a complete set of chromosomes have a higher chance of survival and persistence into the second- and old-growth stages. The strong differences in plant performance for height and volume gains among cultivars reported in [55] might be explained by such competition between aneuploid and euploid clonal sprouts.

While our statistical analysis did not reveal a significant difference between old-growth and second-growth stands, the observation of aneuploidy exclusively in second-growth stands is still noteworthy. As explained above, it is possible that individual second-growth plants might have both euploid and aneuploid tissues, but as the tree grows, the aneuploid tissue is outcompleted by the euploid material. Some evidence to this point is provided in the work of Zane Moore [45] who identified one aneuploid branch in an old-growth tree with the seeds collected from that branch being fully sterile.

Our study did not aim to determine the origins or causes of aneuploidy (meiotic vs. mitotic). Aneuploidy also does not seem to be limited to plants of only clonal origin. Yet, clonal plants do have an advantage as compared to the plants originating from seed because they don’t need to put resources into root system development. After a tree is cut (or damaged, for example after a fire), it can actively resprout and re-establish itself. It is possible that resource availability protects clonal sprouts from the negative effects of dosage imbalance in the early stages of plant development, but as the seed trees become mature, they might outcompete vegetative trees.

### 5.4 Management implications

A common goal in redwood restoration projects is the return of the old-growth forest structure [56–58] after the intensive harvesting of the previous century. However, to meet the old-growth structure objective, it is preferable that genetics of individual ramets are taken into account. It is possible that not all redwood clones might be able to reach the old-growth stages, due to the fitness differences between aneuploid and euploid trees.

Importantly, here we do not advocate for the systematic removal of aneuploid trees in conservation efforts to achieve the goal of old-growth structure. In certain cases, and in some redwood stands it might be best for long-term restoration goals to use multiple restoration strategies, including natural recovery [59]. While aneuploidy in general has negative consequences for the fitness of an individual, it can also provide evolutionary flexibility by promoting genome and chromosome instability (CIN), facilitating cellular adaptation, and redistribution of resources within a cell [60–62]. Additionally, while our results suggest strong fitness effects of aneuploidy, more studies are necessary to quantify those fitness effects, and specifically the phenotypic differences between euploids and aneuploids to undestand the effects of aneuploidy on tree survival, seed viability etc and evaluate the tree performance not only in the sense of the short-term growth but also its ability to survive in the changing climate.

The species is also a valuable timber resource and managing this resource might require different approaches than in restoration. Many of the redwood commercial stands are currently managed on short rotations of approximately 50 years [56], and in such stands persistence of the trees into the old growth states as well as seed viability might not be the main objective. After harvesting, such stands are also restored using the tissue-culture planting material. Pre-commercial thinning of such stands is often recommended to increase the stand volume increment. This is when avoiding planting aneuploids becomes especially important.

Caution should be factored into the decision-making process regarding which cultivars to plant in the field, given the prevalence of aneuploids with a missing chromosome among the tissue-cultured plants. The initial performance of the seedlings should be taken into account, as the negative effects of aneuploidy are likely to manifest themselves in the very early stages of development. Another important factor to take into account is how many generations of the tissue-culture replication a particular clonal line has been through. It is not yet clear how aneuploidy propagates throughout clonal generations, but some examples from the literature indicate that aneuploidy remains and increases in plants propagated by selfing. In *Brassica napus*, for example, aneuploidy increased from 24% to 94% after only 10 generations [63].

Identifying aneuploid trees either in tissue-culture or in the second-growth stands is another outstanding question. There are methods to identify aneuploidy in human cells and it should not be difficult to adapt those methods to identification of aneuploids in plants. Quantitative PCR (qPCR) is a common approach that detects aneuploidy by amplifying DNA sequences from targeted chromosomes and quantifying them in real time, comparing the results against a reference chromosome to determine if extra or missing copies are present. More comprehensive methods include fluorescence in situ hybridization (FISH), where fluorescent probes specific to chromosomes are hybridized to cell nuclei, allowing for direct visualization and counting of chromosomes under a fluorescence microscope. However, many of these methods require chromosome-specific probes, which are not yet available for redwood.

## 6 Conclusions

This study aimed to evaluate instances of aneuploidy in coast redwood, assuming a baseline ploidy of six. Our analysis shows that coding sequence are found in sets of six and they are the most abundant category, reflecting the genome’s hexaploid nature. At the population level, we observe structural instability due to aneuploidy, with extra chromosomes more common than chromosome loss. Unique among forest trees, coast redwood is a rare hexaploid and autopolyploid conifer [16] that reproduces vegetatively and can live for thousands of years, making structure genome errors costly. Aneuploidy is present in second-growth populations, where extra chromosomes are more common than a chromosome loss, whereas tissue culture plants mainly exhibit missing-chromosome aneuploidy. These findings have significant implications for coast redwood restoration and management.

## 8 Acknowledgements

The authors thank undergraduate researchers Liam Galleher, Claire Whicker, Simone Stevens, Nic Dutch, and Jenifer Camarena for their assistance in sample collection and DNA extraction. We also thank Michelle Davila for help in library preparation. We are also grateful to participating land managers who provided access to sampling locations, including those at The Napa Valley Reserve (Paul Asmuth), California State Parks, The California Department of Forestry and Fire Protection(CalFire) and especially Lynn Webb, Green Diamond Resource Company (Carlos Gantz), Mendocino Redwood Company, Humboldt Redwood Company, and The Lyme Timber Company.

## 9 Data availability

The raw sequencing data generated in this study have been deposited in the NCBI Sequence Read Archive (SRA) under the accession number SUB14730245 (BioProject: PRJNA1163354). The data are publicly accessible and can be freely downloaded for academic and research purposes.

## 10 Author contributions

Conceptualization: [ASN, RD, and RN]. Methodology: [ASN, RD, RN, LS, JS]. Data Collection: [ASN]. Data Analysis: [ASN, RN, JS]. Writing – Original Draft: [ASN, RN, LS]. Writing – Review & Editing: [ASN, RD, RN, LS, JS]. Funding Acquisition: [ASN, RD, RN].

## Appendix A. Read depth normalization by the PCR group

## Appendix B. Trace plots for MCMC parameters

